# A non-human primate model of Amyotrophic Lateral Sclerosis

**DOI:** 10.1101/2025.10.27.684759

**Authors:** Rachael H. A. Jones, Luciano Saieva, Fabien Balezeau, Ian Schofield, Caroline McCardle, Mark R. Baker, Stuart N. Baker

**Author notes:** Senior author (Stuart N. Baker).

## Abstract

Approximately 97% of patients with amyotrophic lateral sclerosis (ALS) have cytoplasmic mislocalization and aggregation of the ubiquitous nuclear protein, TDP-43. Current rodent models of this disease fail to replicate the progressive motor weakness and characteristic histopathology, possibly because of fundamental neuroanatomical and genetic differences between rodents and humans. In this study, the TDP-43 protein was overexpressed in the motor neuron pool of the brachioradialis muscle unilaterally in two six-year-old female rhesus macaques, using an intersectional genetics approach involving infection with genetically modified adeno-associated virus. Magnetic resonance images demonstrated delayed signal hyperintensities limited to the injected brachioradialis that persisted for 6-7 weeks, consistent with motor neuron degeneration and denervation of the targeted muscle. At *post-mortem*, the virus-mediated focal protein overexpression event was found to induce widespread deposits of pathological phosphorylated TDP-43 throughout the cervical spinal cord and motor cortex bilaterally, indicating an ALS-like spread of proteinopathy from the transfection site.

## INTRODUCTION

Amyotrophic lateral sclerosis (ALS) is a rapidly progressive adult-onset neurodegenerative disorder characterized by degeneration of both upper motor neurons in the motor cortex (corticospinal cells) and lower motor neurons (α-motor neurons) in the spinal cord and bulbar cranial nerve nuclei[1]. It affects 1-2 in 100,000 people globally[2], with over 90% of cases occurring sporadically in patients with no family history of the disease[3]. Misfolded, protein-rich inclusions are a frequent molecular hallmark of neurodegenerative disorders. In ALS, a proteinopathy involving the 43-kDa trans-active response DNA-binding protein (TDP-43) is found in 97% of cases[4]. TDP-43 becomes mislocalized to the cytoplasm of motor neurons and hyperphosphorylated (pTDP-43), and aggregates to form pathological inclusions[5-7]. These histopathological changes spread to adjacent neural tissue and also to more distant anatomically connected regions[7], at the same time as the evolving patterns of disease observed clinically[8].

The molecular mechanisms of pathological spread between affected cells remain unknown; in ALS a secondary question concerns the direction of disease spread. One hypothesis is that the disease process begins in the motor cortex and spreads from there to bulbar and spinal motor neurons and other cortical areas[9-12]. This corticofugal, or ‘dying forward’ model directly implicates the long corticospinal/corticobulbar axons, and their monosynaptic connections to alpha motor neurons. It is supported by the post-mortem identification of pathological pTDP-43 in both upper and lower motor neuron populations even in the first stage of disease[7], and by the functional sparing of extraocular and sphincter muscles, whose innervating motor neurons do not receive direct cortico-motoneuronal inputs[13, 14]. However, corticofugal spread does not account for rarer ALS phenotypes which affect lower or upper motor neurons selectively (progressive muscular atrophy and primary lateral sclerosis respectively)[15]. Alternate hypotheses, for example that the disease is initiated in peripheral nerve following injury and propagates backwards[16, 17] are reviewed by Baker [18].

Since the identification of causative mutations in the *SOD1* gene[19], and subsequently other genes responsible for monogenic familial amyotrophic lateral sclerosis, rodent models have dominated research into ALS. However, as in patients with *SOD1* mutations, *SOD1* transgenic mice have no TDP-43 proteinopathy and little evidence of upper motor neuron involvement[20, 21]. The rodent motor system has fundamental differences from humans; importantly, it lacks the direct cortico-motoneuronal connections which may be important in disease spread[22]. To address fundamental questions of pathogenesis, and provide a means of testing novel disease-modifying compounds, an animal model that accurately reproduces the progressive motor weakness, characteristic histopathologies, extended pre-symptomatic phase and subsequent rapid deterioration of human ALS is urgently required.

One previously published approach used adeno-associated virus (AAV) mediated TDP-43 overexpression in the spinal cord to recapitulate the ALS phenotype in adult macaque monkeys[23]. These animals displayed progressive muscle weakness accompanied by cytoplasmic pTDP-43 inclusions in post-mortem tissue. However, because viral transfection and gene overexpression was widespread throughout the spinal cord, it was not possible to address questions of disease spreading. Here we have built on this previous work, using an intersectional genetics approach to overexpress wild-type human TDP-43 in a focal subset of spinal motor neurons in macaques. This study was based in part upon the first author’s doctoral thesis[24]. Using magnetic resonance (MR) imaging, we detected regions of signal hyperintensities, which developed exclusively in the injected brachioradialis muscle and persisted for six to seven weeks in each animal. These bands of elevated signal, which were detected eight and fourteen weeks after virus injection in each animal respectively, are consistent with signal changes previously noted in denervated muscle[25-27] and likely indicate the onset of TDP-43 toxicity resulting from TDP-43 overexpression within the transfected spinal motor neurons. In this way, muscle signal increases on MRI could be used as a marker for the onset of virus mediated pathological change. Animals were terminated seven months after viral transfection, and cervical motor neurons and large pyramidal cells of the primary motor cortex examined histologically. Although initial overexpression of TDP-43 was limited to the brachioradialis motor neuron pool, at *post mortem* we found widespread deposits of pathological phosphorylated TDP-43 throughout the cervical spinal cord and motor cortex bilaterally. The absence of fluorescent reporter protein, which had been included in the viral genetic payload, together with a reduction in motor neuron counts within spinal segments innervating brachioradialis, was consistent with complete degeneration of the initially transfected motor neurons. These results demonstrate that focal overexpression of TDP-43 in a spinal motor pool is sufficient to induce widespread pathological TDP-43 inclusions throughout the spinal cord and motor cortex. We expect that this animal model will allow novel disease-modifying agents to be tested, with a high chance of results successfully translating to human patients.

## STAR★METHODS

### KEY RESOURCES TABLE

| REAGENT OR RESOURCE | SOURCE | IDENTIFIER |
| --- | --- | --- |
| Antibodies |  |  |
| RFP Pre-adsorbed Polyclonal Antibody | Rockland | Cat#600-401-379;<br>RRID:AB_2209751 |
| Goat Anti-Choline Acetyltransferase polyclonal antibody | Sigma-Aldrich | Cat#ab144P;<br>RRID:AB_2079751 |
| TDP-43 Polyclonal antibody | ProteinTech | Cat#10782-2-AP;<br>RRID:AB_615042 |
| Anti TDP-43, phospho-Ser409/410 mAb (Clone 11-9) | CosmoBio Ltd | Cat#CAC-TIP-PTD-M01;<br>RRID:AB_1961900 |
| Donkey Anti-Rabbit IgG H&L (Alexa Fluor® 594) Antibody | abcam | Cat#ab150076;<br>RRID:AB_2782993 |
| Donkey Anti-Goat IgG H&L (Alexa Fluor 488) Antibody | abcam | Cat#ab150129;<br>RRID:AB_2687506 |
| Bacterial and virus strains |  |  |
| AAV9-EF1a-FLEX-FLAG_TDP-43-WPRE | This paper | N/A |
| rAAV2-retro-pAAV-hSyn-CRE-tdTomato | This paper | N/A |
| Biological samples |  |  |
| Healthy Macaca Mulatta brain and spinal cord tissue | MRC Centre for Macaques | N/A |
| Chemicals, peptides, and recombinant proteins |  |  |
| Tween-20 | Sigma-Aldrich | Cat#P1379; CAS: 9005-64-5 |
| Donkey Serum | Bio-Rad Laboratories | Cat#C06SB |
| Sudan Black B | Sigma-Aldrich | Cat#199664; CAS: 4197-25-5 |
| Fluoroshield mounting medium with DAPI | Abcam | Cat#ab104139 |
| Histo-clear II | Scientific Laboratory Supplies Ltd | Cat#NAT1334 |
| VectaMount® Express Mounting Medium | Vector Laboratories | Cat#H-5700-60 |
| Critical commercial assays |  |  |
| ImmPRESS HRP goat anti-rabbit IgG polymer reagent | Vector Laboratories | Cat#MP-7451 |
| DAB Substrate Kit, Peroxidase (HRP), with Nickel, (3,3'-diaminobenzidine) | Vector Laboratories | Cat#SK-4100 |
| Deposited data |  |  |
| Raw and analyzed data | This paper | https://data.ncl.ac.uk |
| Experimental models: Organisms/strains |  |  |
| Macaca Mulatta wild-type | MRC Centre for Macaques | N/A |
| Software and algorithms |  |  |
| ImageJ | Schneider et al.[28] | https://imagej.nih.gov/ij/ |
| MRtrix3 | Tournier et al.[29] | https://www.mrtrix.org/ |

## METHOD DETAILS

### ANIMALS

The main study was performed on two adult female monkeys (*Macaca mulatta*; Monkeys Mi and Ma, age six years, weight 6.1 and 6.9kg at outset). Viral vector efficacy studies were performed in 5 adult female macaques (Monkeys Wa, Wi, Y, Sd, Sp; ages four to five). In addition, control spinal cord tissue was collected post-mortem from one female macaque (Monkey Z, age 3 years), and control M1 tissue from another female macaque (Monkey W, age 3 years). Neither had been involved in electrophysiological recordings, and were housed at the MRC Centre for Macaques. All animal procedures were performed under appropriate licenses issued by the UK Home Office in accordance with the Animals (Scientific Procedures) Act (1986) and were approved by the Animal Welfare and Ethical Review Board of Newcastle University.

### SURGICAL PREPARATION

During the MRI acclimatization phase (see below), an MRI scan of the head was performed under general anesthesia. This was used to design an annular headpiece with underside shaped to fit the skull; the headpiece was manufactured from MRI-compatible plastic (TECAPEEK MT CF30, Ensinger, Nufringen, Germany) using a computer-controlled milling machine.

After MRI acclimatization was complete, both monkeys underwent a sterile implant surgery. After initial sedation with ketamine (10mg·kg^-1^ IM), anesthesia was induced with medetomidine (3 μg·kg^−1^ IM) and midazolam (0.3mg·kg^-1^ IM). The animal was intubated, and anesthesia maintained using inhalation of sevoflurane (2.5-3.5% in 100% O_2_) and IV infusion of alfentanil (0.4 μg·kg^−1^·min^−1^). Methylprednisolone was infused to reduce oedema (5.4 mg·kg^−1^·hr^-1^ IV). Blood-oxygen saturation, heart rate, arterial blood pressure (using a non-invasive blood pressure cuff on the leg), core and peripheral temperature and end-tidal CO_2_ were monitored throughout; ventilation was supported with a positive pressure ventilator. Hartmann’s solution was infused to prevent dehydration (total infusion rate including drug solutions 5–10 ml·kg^−1^·h^−1^). Body temperature was maintained at 37°C using a thermostatically controlled electric heating blanket placed under the animal and a warm air blanket (3M Bair Hugger, Ontario, Canada) on top of the animal. Intraoperative prophylactic antibiotics (cefotaxime 20mg·kg^-1^ IV) and analgesia (carprofen 5 mg·kg^-1^ SC) were given. The headpiece was fixed to the skull using ceramic bone screws (Thomas Recording, Giessen, Germany) and dental acrylic. In the same surgery, fine wire EMG electrodes were implanted in eight muscles in each forelimb, and nerve cuffs placed on the median and radial nerves bilaterally; wires from these electrodes were tunneled to connectors on the headpiece. Data from EMG and nerve cuff implants are not presented in this publication. Post-operative care included a full program of antibiotics (co-amoxiclav, dose as above) and analgesics (meloxicam, 0.2mg·kg^-1^ oral plus a single dose of buprenorphine 0.02mg·kg^-1^ IM).

### MRI SCANS OF THE FOREARM

Animals were acclimatized to undergo MRI scans of the forearm in the awake state, using positive reinforcement training. A laser scanner (Go!Scan, Creaform 3D, Levis, Quebec, Canada) was used to create a digital model of the forearm, from which a rigid plastic cast was manufactured by 3D printing. This individualized cast slid into a cylindrical attachment on the door of the primate chair, fixing the elbow in 90° flexion, with the forearm semi-pronated so that the radius and ulna were oriented in a vertical plane. Within the attachment was a custom transmit/receive radiofrequency coil, positioned directly over the region of interest in the forearm. This allowed regular MRI scanning with the upper limb held in a consistent position throughout and between sessions. A camera on the MRI chair allowed behavior to be monitored constantly; the monkey was given fruit juice rewards for sitting quietly while in the bore of the scanner via a pump system operated by the investigator. After the headpiece implant (see above), the monkeys were also trained to accept atraumatic head-fixation during scans to reduce motion artefact. Both right and left arms were scanned on different days in each week, allowing progressive changes to be tracked.

MRI scans were acquired using a vertical 4.7 Tesla research MRI scanner (Bruker BioSpin, Ettlingen, Germany). T2-weighted Rapid Acquisition with Relaxation Enhancement (RARE) scans were used. A fat-selective radiofrequency pulse was used to reduce signal artefacts produced by fat in the subcutaneous fat layer, bone marrow and surrounding the muscles of the forearm. Repetition time (TR) was 7900ms, and echo time (TE) was 57.3ms. The scans were acquired as axial slices with a resolution of 0.55mm, a 128×128 in-plane matrix, and 64 slices with a slice thickness of 1.88mm. Eight scans were acquired in 18min and combined with median averaging to improve the signal-to-noise ratio and reduce motion artefacts.

### MRI POST PROCESSING

Raw scans were pre-processed, segmented and filtered (median and Gaussian) in Fiji ImageJ (NIH)[28]. Each scan was then realigned and registered using the mrregister function on MRtrix3[29]. Scans from each session were ordered by date and concatenated. Each image was normalized against a Gaussian-filtered region of interest (ROI) measured in the flexor compartment, an area not expected or shown to display virus-related signal change. For all slices along the proximo-distal axis (Monkey Mi: 34 slices; Monkey Ma: 40 slices), the brachioradialis muscle was demarcated with a hand-drawn contour within which signal intensity measurements were taken. Mean signal intensity was computed by averaging all voxels within the obtained ROI. Intramuscular blood vessels were masked to remove signal artefacts. Both left and right arm scans were analyzed using identical methodology.

To compare between sessions, the mean signal intensity (*S*) from each session (*N*) was normalized by subtracting the mean signal intensity in the baseline sessions 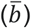, and dividing by the standard deviation of the baseline *σ*_*b*_ to give a *t* score for each session, as described in Equation (1):

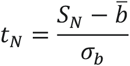

Data were analyzed using custom scripts written in the MATLAB environment (R2020a, MathWorks). Mean signal intensity in the proximal to distal z slices was smoothed with a moving window of five z slices. Z slice windows were tested against their respective baseline sessions. A total of nine scans spanning sessions prior to and immediately following the virus injections contributed towards the baseline period for the left and right arms of both animals. All statistical tests were performed using custom MATLAB scripts. The Benjamini-Hochberg procedure was performed to correct for multiple comparisons[30].

### VIRUS INJECTIONS

The objective of the viral transfection was to overexpress TDP-43 in motor neurons projecting to a single muscle, the brachioradialis. We performed pilot experiments in 5 adult female macaques (Monkey Wa, Wi, Y, Sd, Sp) as part of different studies, to test different viruses, serotypes and viral titers (Fig. 4A). Viral transfection was successful in two of these pilot animals, however efficacy was low, with around one motor neuron expressing detectable fluorescent tag (tdTomato) per 40µm section. We therefore chose to use an intersectional approach for our main study, in which a flipped, antisense copy of the TDP-43 gene was introduced into many motor neurons throughout the spinal cord. A second virus was delivered to a single motor neuron pool selectively by retrograde transport from the muscle. This inverted the TDP-43 gene, allowing it to be selectively expressed (Fig. 1).

**Figure 1.**
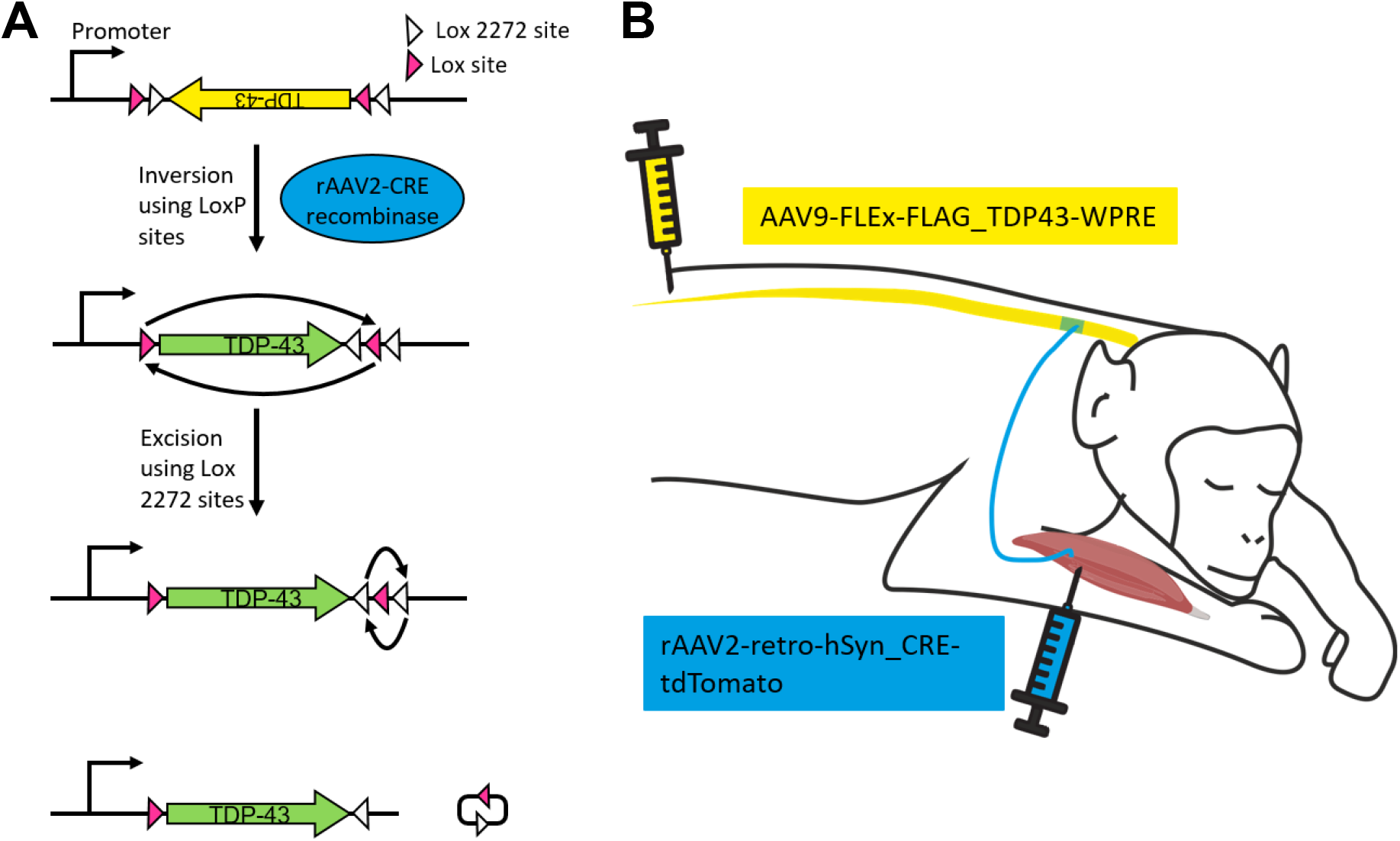
Intersectional genetics approach to overexpress TDP-43 in a single spinal motor neuron pool of two rhesus macaques. (**A**) The FLEx system exploits Cre recombinase to trigger the inversion and excision of the gene of interest from between two loxP sites, thus ‘flipping’ the gene into an active configuration. (**B**) Inactive human wild-type TDP-43 DNA packaged in an AAV9 vector was injected intrathecally via lumbar puncture. The animals were placed in the Trendelenburg position for 10 minutes after this injection to facilitate rostral spread of the virus to the cervical segments [31]. In the same procedure, a second virus, rAAV2 retro, was injected into the right brachioradialis muscle. This vector encoded Cre recombinase. The TDP-43 gene will be switched into the active, readable state in any cells expressing both viruses. This system ensured that transduction was limited to the motor neurons innervating the right brachioradialis muscle.

While anesthetized and under sterile conditions, Monkeys Mi and Ma underwent a lumbar puncture in which a 21G hypodermic needle was inserted between the fourth and fifth lumbar spinous processes, identified by palpation of spinal landmarks. An intrathecal injection through this needle was made of AAV9-EF1a-FLEX-FLAG_TDP-43-WPRE (5.71 × 10^13^vg/ml, volume injected 1ml). This injection was followed by an epidural blood patch to minimize the risk of CSF leak. The animal was moved from the lateral decubitus position into the Trendelenburg position, in which the animal lay supine, head downwards at a 15-30° angle for 10 minutes. This position has been shown to improve transduction efficacy of cervical motor neurons following intrathecal administration of AAVs [31]. While in this position, the animal was injected with rAAV2-retro-pAAV-hSyn-CRE-tdTomato (8.96 × 10^12^ vg/ml in 500μl) into five sites in the right brachioradialis muscle (Fig. 1). The injection needle was insulated apart from its tip; injections were made at sites with low threshold for eliciting a twitch after electrical stimulation through the needle, presumed to correspond to the motor end plates (biphasic pulses, 0.2ms per phase using isolated constant current stimulator, AM Systems model 2200). Twitch thresholds were 100-250μA for monkey Mi, and 50-150μA for monkey Ma. After this procedure, the non-steroidal anti-inflammatory drug meloxicam was given (0.2 mg/kg, immediately after the procedure and once on the following day).

### TISSUE PREPARATION FOR HISTOLOGY

Following the completion of the study (224 days after virus injection for both monkeys Mi and Ma), the monkeys were killed by overdose of anesthetic (propofol IV) and perfused through the heart with phosphate buffered saline (PBS) followed by 10% neutral buffered formalin. The cortex, cerebellum, brainstem and spinal cord were removed and immersed in 10% neutral buffered formalin for 16 hours at 4°C before progressing through ascending concentrations of PBS sucrose (10%, 20%, 30%) for cryoprotection. Cortical blocks containing the mediolateral extent of the left and right sensorimotor cortex were removed. The spinal cord from C1-T1 was separated into blocks. A freezing sledge microtome (8000-01, Bright Instruments Co. Ltd, United Kingdom) connected to a solid-state freezer (53024-01, also Bright Instruments Co. Ltd) was used to cut the spinal cord segments into 40µm coronal sections and cortical blocks into 60µm parasagittal sections. All sections were stored free floating in PBS at 4°C until further processing.

Tissue underwent either fluorescent immunohistochemistry to detect the fluorescent viral tag, tdTomato, and for the identification and counting of motor neurons with choline acetyl transferase (ChAT) or DAB immunohistochemistry for TDP-43 and phosphorylated TDP-43. Brightfield images of TDP-43 and pTDP-43 labelled sections were acquired using a Zeiss AxioImager (Carl Zeiss AG, Oberkochen, Germany), which allowed for tiling using a 10x objective. Fluorescently labelled spinal cord sections were imaged using a Zeiss CellDiscoverer 7 (Carl Zeiss AG, Oberkochen, Germany) connected to a highly sensitive Hamamatsu Fusion sCMOS camera, which allowed for multi-slide imaging. Full details of histological protocols are available in the Supplementary Methods.

### IMMUNOHISTOCHEMISTRY TO DETECT FLUORESCENT VIRAL TAG

This procedure aimed to detect the tdTomato fluorescent protein, which was encoded in the genetic payload of the rAAV2-retro virus. Sections were collected at 960µm intervals from segments C4-C8 (known to supply the brachioradialis muscle) and labelled with an antibody against red fluorescent protein and its variants (600-401-379, RFP Pre-adsorbed Polyclonal Antibody, Rockland; 1:1000). Primary antibody dilutions were performed using a PBS solution containing 0.3% Tween-20 (Sigma-Aldrich) and 5% donkey serum (Bio-Rad Laboratories). The primary antibody solution was applied to the sections overnight at 4°C. The sections were washed in PBS three times before donkey anti-rabbit Alexa Fluor 594 (ab150076, abcam; 1:1000) was applied as a secondary antibody for two hours at room temperature. The sections were shielded from light during the secondary incubation and for all subsequent steps to prevent bleaching of the signal. The sections were washed three times in PBS before being mounted onto slides (Superfrost Plus, Thermo Scientific). Once the sections had dried and adhered onto the slides, they were dehydrated in 70% ethanol for five minutes and immersed in Sudan Black B (199664, Sigma-Aldrich) dissolved in a 70% ethanol solution for 10 minutes. Following this the slides were washed twice in PBS to remove any residual Sudan Black deposits. Sudan Black binds to and quenches the autofluorescent protein lipofuscin, which builds up in the cytoplasm of cells of the CNS with age[32]. Fluoroshield mounting medium with DAPI (4′,6-diamidino-2-phenylindole) (ab104139, Abcam) was used to counterstain cell nuclei and coverslips were placed.

### MOTOR NEURON LABELLING

To allow quantitative counts of motor neuron number, spinal cord sections were collected at 480-720µm intervals and labelled with a primary antibody for choline acetyl transferase (ChAT) (ab144P, Sigma-Aldrich; 1:500). Primary antibody dilutions were performed using a PBS solution containing 0.3% Tween-20 (Sigma-Aldrich) and 5% donkey serum (Bio-Rad Laboratories). A goat anti-ChAT polyclonal antibody (Sigma-Aldrich; 1:500) was applied to the sections overnight at 4°C. The sections were washed in PBS three times before donkey anti-goat Alexa Fluor 488 (ab150129, abcam; 1:1000) was applied as a secondary antibody for two hours at room temperature. Subsequent steps were repeated as above.

### LABELLING OF TDP-43

Spinal cord sections were collected at 960µm intervals, and cortical sections were collected at 1440µm intervals for TDP-43 immunostaining (10782-2-AP, TDP-43 Polyclonal antibody, ProteinTech; 1:2000). Further spinal cord sections were collected at 320-480µm (Mi = 1/12, Ma = 1/8) intervals, and sensorimotor cortex sections were collected at 720µm intervals, for phosphorylated TDP-43 (pTDP-43) immunostaining (CAC-TIP-PTD-M01, Anti TDP-43, phospho-Ser409/410 mAb (Clone 11-9), CosmoBio Ltd; 1:500)[33]. Sections were washed with 0.1M PBS before being incubated with 3% H_2_0_2_ for five minutes to quench any endogenous peroxidase activity. Following this, the sections were washed with PBS three times. Primary antibody dilutions were performed using a PBS solution containing 0.3% Tween-20 (Sigma-Aldrich) and 5% goat serum (Vector Laboratories). A rabbit anti-TDP-43 polyclonal antibody (ProteinTech; 1:2000) was applied to the sections for TDP-43 analysis overnight at 4°C, while sections for pTDP-43 processing were incubated with a mouse anti-TDP-43 phospho-Ser409/410 polyclonal antibody (CosmoBio Ltd; 1:500) overnight at 4°C. The following day the sections were washed three times with PBS prior to a 30-minute incubation with an ImmPRESS HRP goat anti-rabbit IgG polymer reagent (MP-7451, Vector Laboratories) at room temperature. The sections were again washed with PBS 3 times to remove any residual secondary antibody. A DAB substrate (3,3’-diaminobenzidine) (SK-4100, Vector Laboratories) was applied to the sections until a desired stain intensity developed (two minutes). The DAB substrate produces a brown precipitate in the presence of the peroxidase (HRP) enzyme. Tap water was applied to the sections for five minutes to quench the reaction. At this point sections were mounted onto slides and allowed to air dry. Sections were dehydrated in ascending ethanol solutions (70%, 90%, 100%) and cleared in histoclear (Sigma-Aldrich) for 10 minutes. Sections were coverslipped with VectaMount (Vector Laboratories).

### MICROSCOPY AND QUANTIFICATION

Spinal cord motor neurons were identified by their morphology, as large multipolar neurons, and location within lamina IX of the ventral horn grey matter. Non-motoneuronal cell types were excluded from counts based on these criteria. Brightfield images of TDP-43 and pTDP-43 labelled sections were acquired using a Zeiss AxioImager (Carl Zeiss AG, Oberkochen, Germany), which allowed for tiling using a 10x objective. Fluorescently labelled sections were imaged using a Zeiss CellDiscoverer 7 (Carl Zeiss AG, Oberkochen, Germany) connected to a highly sensitive Hamamatsu Fusion sCMOS camera which allowed for multi-slide imaging. Motor neurons labelled for pTDP-43 or ChAT were counted using the ImageJ Cell Counter tool[34] in the left and right ventral horn of all cervical segments for both monkeys.

Betz cells were identified based on their location in layer V of the motor cortex and their large pyramidal cell body. All brightfield and widefield fluorescence images were acquired using a Zeiss AxioImager (Carl Zeiss AG, Oberkochen, Germany), which allowed for the capture of high magnification images as well as tiling using a 10x objective. For analysis, the primary motor cortex was divided into Old M1 and New M1 based on previously described anatomical landmarks[35, 36]. The presence of the giant pyramidal Betz cells was used to demarcate the most rostral border of Old M1 and most caudal border of New M1 for each section analyzed. Betz cells positive for pTDP-43 immunoreactivity within these regions were counted in every 12^th^ section throughout the mediolateral extent of the motor cortex (every 720µm) using the ImageJ Cell Counter tool[34]. The number of counted cells were averaged for each area. To compensate for differences in the size of Old M1 and New M1, the density of pTDP-43 positive cells was measured. Cell density was calculated by dividing the number of counted cells per section by the rostro-caudal length of layer Vb, measured in ImageJ. Before the length could be measured, each image was spatially calibrated to convert the number of pixels into a real distance in micrometers.

### STATISTICAL ANALYSIS

Data were analyzed using custom scripts written in the MATLAB environment (R2021a, MathWorks). Statistical tests were performed using MATLAB. Paired t-tests were used to compare pTDP-43 and ChAT positive cell counts from contralateral sides of each spinal segment. A two-way ANOVA was used to compare both the effects of side (ipsilateral-contralateral) and the effect of segment on pTDP-43 cell counts. To examine intersegmental differences, pairwise t-tests were performed to compare each segment against the group of segments in which significant reductions in ChAT+ motor neurons were counted. Unpaired t-tests were used to compare counts from different hemispheres and between cortical subdivisions. Benjamini-Hochberg corrections were applied to all statistical tests when necessary to account for multiple comparisons[30], with a significance level of p<0.05; in the text, uncorrected p values are given, together with a statement of whether these should be considered significant after this correction.

## RESULTS

### MRI CAN DETECT THE ONSET OF VIRUS-MEDIATED DENERVATION IN BRACHIORADIALIS MUSCLE

RARE-MRI scans of equivalent axial slices of the left and right forearms of both monkeys are shown in Figure 2, at different time points relative to virus injection. Signal intensity appeared constant throughout the muscles of the forearm, including the brachioradialis muscle (delineated in red in Fig. 2) prior to the virus injections, except for a mild hyperintensity throughout the extensor compartment in the right forearm of Monkey Mi. This may relate to earlier slight damage to the radial nerve in this arm, caused during the implantation of a nerve cuff. This diffuse hyperintense signal had normalized by 6 weeks after the virus injection, when the signal was uniform in all forearms scanned (Fig. 2C, D).

**Figure 2.**
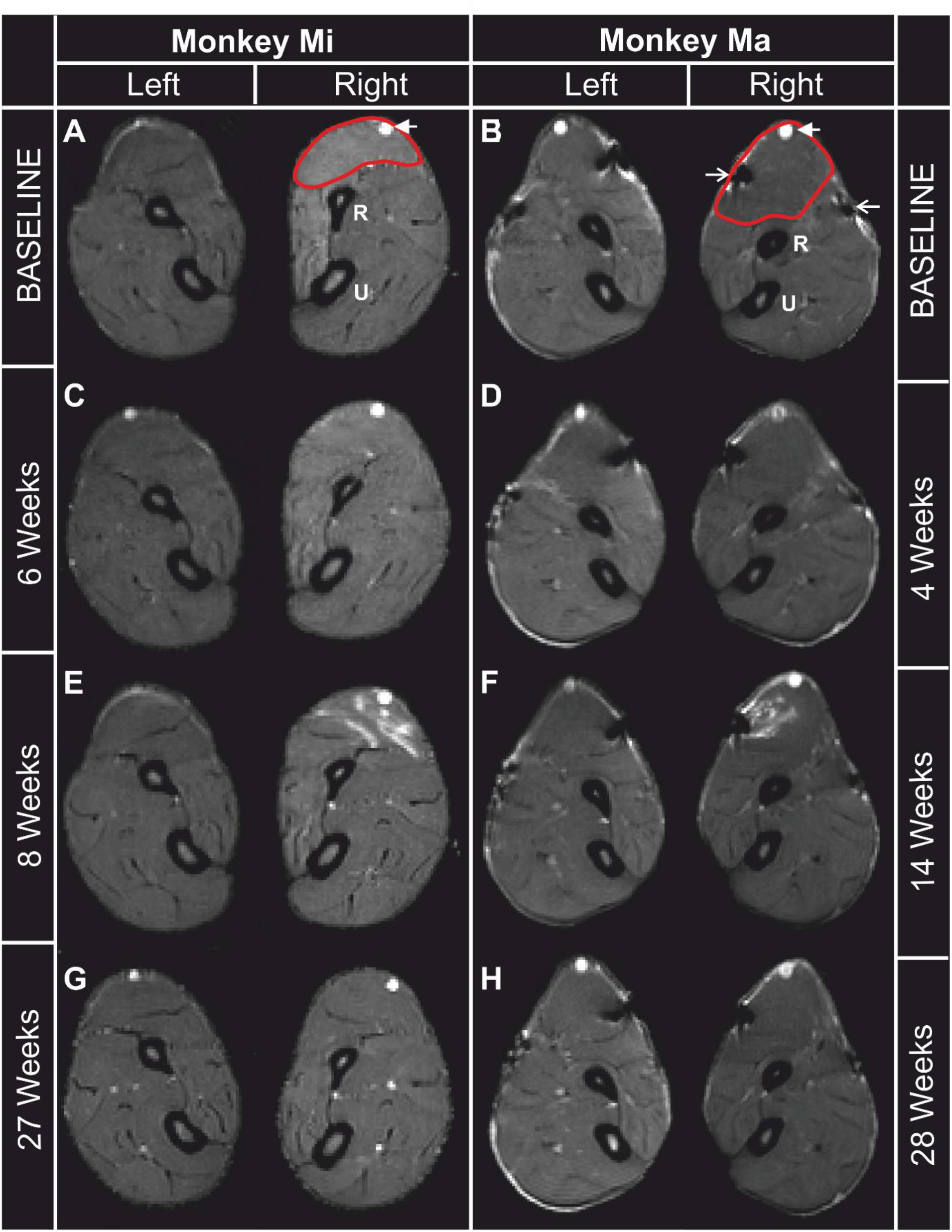
Muscle signal intensity from T2-weighted Rapid Acquisition with Relaxation Enhancement (RARE) scans. All panels show cross-sectional images of the left and right arms from two monkeys. For orientation, on a single scan for each animal the brachioradialis muscle is demarcated with a red boundary, the ulnar (U) and radius (R) bones are labelled, and a superficial blood vessel indicated with the filled arrowhead. In some scans, artifacts caused by the implanted EMG wires can be seen in both a flexor and extensor muscle of monkey Ma, indicated with an open arrowhead. (**A, B**) Baseline scans prior to virus injection. (C, D) 4-6 weeks after virus injection, no signal abnormalities are seen. (**E, F**) 8-14 weeks after virus injection, bands of hyperintense signal are seen localized to the brachioradialis muscle. (**G, H**) Signal abnormalities normalize after 8 weeks in both animals

At 8 weeks (monkey Mi) and 14 weeks (monkey Ma) after viral transfection, bands of high signal began to appear localized to the brachioradialis muscle (Fig. 2E, F). These continued to increase before again normalizing (Fig. 2G, H). None of the other muscles scanned displayed this abnormal signal pattern.

To quantify the time course of these changes in MRI signal, hand-drawn boundaries were used to demarcate the brachioradialis in every proximo-distal axial slice along the muscle (example red outlines in Fig. 2A, B). Mean signal intensity was calculated within this region, and then normalized to the baseline to give a t score as described in Methods. On the left (non-injected) side, there were no significant changes for any proximo-distal location along the muscle (different lines on Fig. 3A, C). On the right (injected) side, the mean signal intensity increased significantly above baseline 9 and 15 weeks after viral transfection in Monkeys Mi (*P*=5.2×10^−7^, Fig 3B) and Ma (*P*=3.4×10^−4^, Fig. 3D), and remained significantly elevated in at least one location for 6 and 7 weeks respectively.

**Figure 3:**
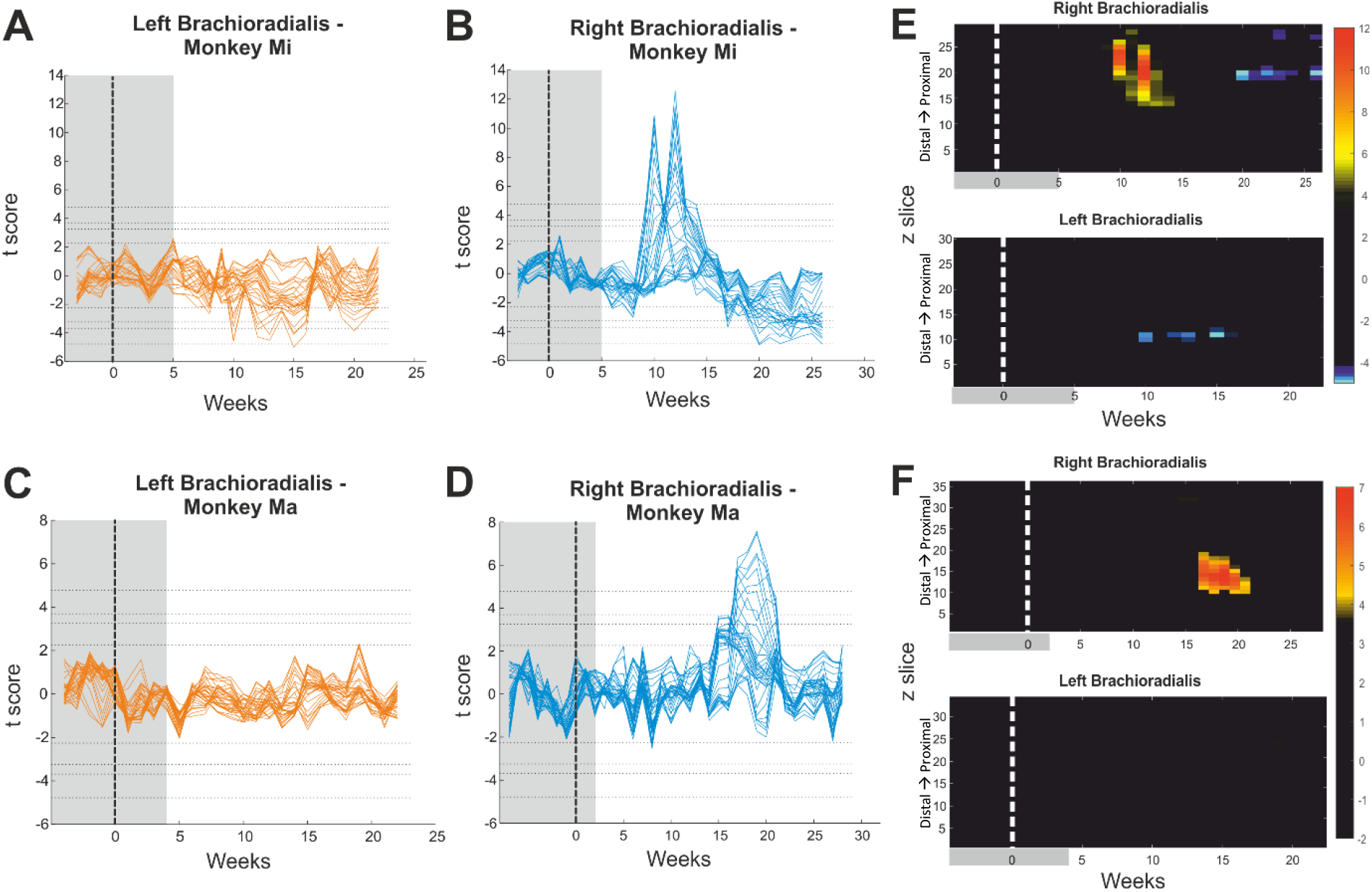
Quantification of RARE-MRI signal intensity changes in the brachioradialis muscle. **(A-D)** Change in signal intensity with time after the virus injection (time 0, vertical dashed line), for left and right sides in two monkeys. Intensity has been normalized by subtracting the mean and dividing by the standard deviation of a baseline period (grey shading). Each colored line presents data from a different proximo-distal slice through the forearm. Horizontal dotted lines mark the significance limits of P<0.05, 0.01, 0.005 and 0.001 after correction for multiple comparisons by the Benjamini-Hochberg procedure. Note the peaks in signal seen on the right side in both animals. (**E, F**) The same data as (A-D), replotted as color maps to show the spatiotemporal pattern. The color scale has been chosen such that black indicates non-significant changes, while red and blue represent significant increases and decreases, respectively. Significance was determined at a threshold of P<0.0017 after correcting for multiple comparisons. Both monkeys showed a focal, time-limited increase in signal in the right BR; in monkey Mi (C) this was followed by a signal decrease. A signal decrease was also recorded during this time in the left BR of monkey Mi. No signal decreases were recorded in the left or right BR of monkey Ma (D).

**Figure 4.**
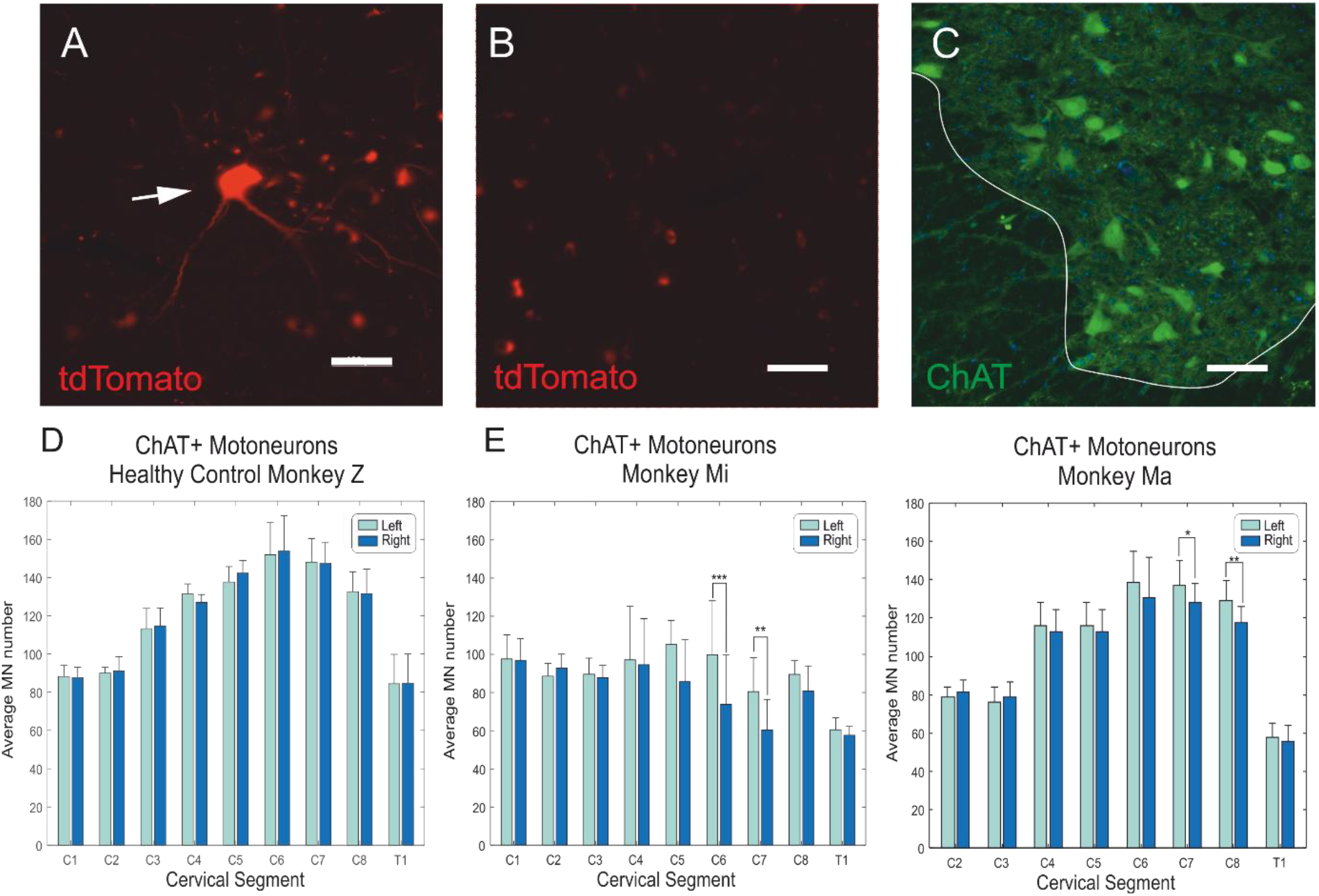
Evidence for motor neuron degeneration in spinal segments innervating the brachioradialis muscle. (**A**) Image from a previous study in our group (Monkey W) involving retrograde viral transfection with rAAV2retro with a tdTomato genetic payload. Motor neurons labelled with the fluorescent protein could be clearly seen. (**B**) Example image from monkey Mi, showing that no motor neurons with tdTomato could be seen. (**C**) Image of ventral horn of the spinal cord from monkey Mi, stained for choline acetyl transferase (ChAT) to identify motor neurons. Scale bar for (A-C) = 200µm. (**D**) Motor neuron counts per section in a healthy control macaque (Monkey Z). There were no significant differences in cell numbers between left and right sides in any segment between C1 and T1. (**E**) Motor neuron counts for the two monkeys infected with TDP-43 constructs as in Figure 1. Asterisks mark significant differences between left and right side (*, P<0.05; **, P<0.01; ***, P<0.0001). Bars are the mean of 6-18 sections; error bars indicate standard errors of the mean.

Figure 3E, F illustrates the changes as a color scale, with the *y* axis representing the distal to proximal location within the brachioradialis muscle, and the x axis time relative to the virus injection (dotted line). In monkey Mi, the significant increase in signal intensity started in a proximal region of the muscle and then moved more distally. In monkey Ma, an initial increase over a wide proximo-distal range became larger and focused more distally.

The results from the right brachioradialis muscle in Monkey Mi showed not only an increase in signal intensity, but a later decrease in signal intensity (*P*=0.0009, Fig. 3B, E). A decrease in mean signal intensity was also evident in the left brachioradialis of Monkey Mi ten weeks after virus injections, 7 weeks earlier than the reduction in right brachioradialis signal intensity (*P*=0.00072, Fig. 3A, E).

### LOSS OF BRACHIORADIALIS MOTOR NEURONS ON THE INJECTED SIDE

The fluorophore tdTomato is a commonly used component of the genetic payload included in AAV viruses, as it reliably allows transfected cells to be identified. In a preliminary experiment using a rAAV2-retro with the same fluorophore, labelled motor neurons could be detected (Fig. 4A, monkey W). However, examination of 26 sections throughout the C4-C8 segments in both monkey Mi and Ma revealed no motor neurons expressing tdTomato (Fig. 4B). We hypothesized that this may be because all transfected motor neurons had degenerated by the time the tissue was examined, seven months after the viral injection. To test for this, 12-18 sections per spinal segment were stained for ChAT (Fig. 4C), and the number of motor neurons on each side counted.

Figure 4D presents the mean number of motor neurons per section from a healthy animal (Monkey Z), which had not undergone viral transfection. Cell counts rose to a maximum in segment C6, corresponding to the cervical enlargement[37, 38]. There was no significant difference between counts on the left and right side at any segmental level (P>0.05). By contrast, in both animals with virus injections there were significantly fewer motor neurons on the right (injected) side than on the left (Fig. 4E), in segments which innervate the brachioradialis muscle[39] (Monkey Mi: C6, *P*=7.8×10^−5^; C7, *P*=0.0062; Monkey Ma, C7, *P*=0.0108; C8, *P*=0.00074; all other comparisons not significant). These results are consistent with focal degeneration of brachioradialis motor neurons transfected by the virus, which would explain the lack of tdTomato fluorophore.

### TDP-43 HISTOPATHOLOGY IN SPINAL CORD

Figure 5A presents example results from cervical spinal cord tissue stained against an antibody for the physiological TDP-43 protein, in the healthy control animal without viral transfection (monkey Z). As previously reported in healthy tissue, dense TDP-43 staining was confined to the nucleus of motor neurons. By contrast, in both animals with viral transfection, TDP-43 was also present in the motor neuron cytoplasm, where it was seen uniformly throughout the cell body and dendrites (Fig. 5B-D). The most noticeable pathology was in the segments innervating the brachioradialis muscle, where a significant reduction in ChAT-expressing motor neurons had been seen (Fig. 4). Some of these cells showed a loss of clear nuclear protein expression alongside substantial dense aggregates throughout the cytoplasm. However, changes were not confined to these segments, but were widespread (Fig. 5C) and included smaller interneurons as well as motor neurons.

**Figure 5.**
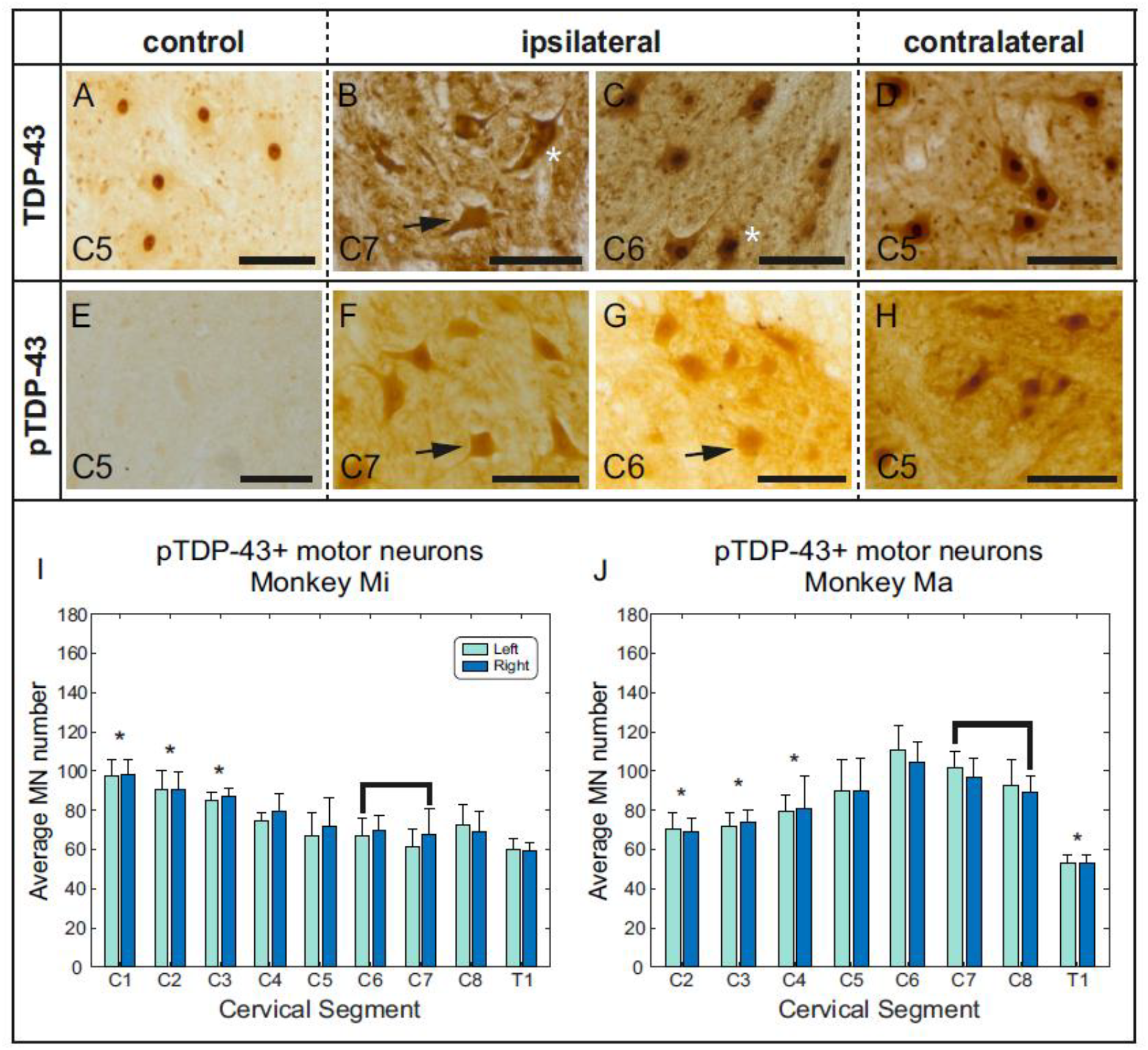
Spinal distribution of TDP-43 and phosphorylated TDP-43 (pTDP-43). (**A-D**) Enlarged view of spinal motor neurons within lamina IX stained for TDP-43 in brown, using DAB. (**A**), from a control animal without virus transfection (monkey Z); TDP-43 was restricted to the nucleus of motor neurons. (**B**) from the C7 segment of Monkey Mi. (**C**) from C6 segment of Monkey Ma. (**D**) from C5 segment of Monkey Mi. Note dense staining in the cytoplasm, with visible clumps of TDP-43, indicated by asterisks (*), and the less prominent nuclear staining in some cells (arrows) in segments which innervate the brachioradialis muscle. (**E-H**) Enlarged view of spinal motor neurons stained for pTDP-43(pS409/410). (**E**) no pTDP-43 staining was identified in a control macaque (monkey Z) without virus transfection. (**F**) from C7 segment of Monkey Ma. (**G**) C6 segment of Monkey Mi. (**H**) C5 of Monkey Ma. Nuclear staining was less prominent in motor neurons of segments innervating the brachioradialis muscle (arrows), than more distant segments. (**I**,**J**) Bar charts showing the counts of pTDP-43+ motor neurons in each spinal segment on the left and right side. There were no significant differences between sides. Asterisks mark segments with significantly different cell counts from the segments which had reduced ChAT+ motor neuron counts. Bars show the mean over 6-18 sections; error bars show the standard error of the mean.

To examine this neuropathology in more detail, we stained the spinal tissue for an antibody against the pathological phosphorylated TDP-43 serine groups 409/410 (pTDP-43). In the control macaque without viral transfection, only very light staining was seen (Fig. 5E). This is most likely attributable to lipofuscin, a lipid protein which accumulates in neurons during normal ageing[32, 40], and is compatible with a complete absence of pTDP-43[33]. In the transfected animals, pTDP-43 was detected in the cytoplasm of motor neurons and smaller dorsal horn and intermediate zone cells, throughout all cervical segments bilaterally (Fig. 5F-H). Nuclear expression of pTDP-43 was also identified. However, this was less pronounced in segments innervating the brachioradialis muscle (Fig. 5H).

Quantification of these results was achieved by counting pTDP-43+ motor neurons in each spinal segment for both transfected monkeys (Fig. 5I, J). A two-way ANOVA confirmed no effect of side (Mi *P*=0.191; Ma *P*=0.107) but revealed an effect of segment (Mi, *P*=3.1×10^−6^; Ma, *P*=7.5×10^−7^). To elucidate the intersegmental effects, pairwise t-tests were performed, comparing each segment to the two segments from each animal with significantly reduced ChAT positive motor neurons (Mi: C6-C7; Ma: C7-C8; Fig. 4E). These *post hoc* tests revealed significant differences between the segments innervating brachioradialis and C1-C3 in Monkey Mi (C1 *P*=2.1×10^−5^; C2 *P*=1.8×10^−4^; C3 *P*=7.4×10^−4^) and C2-C4 and T1 in Monkey Ma (C2 *P*=4.7×10^−4^; C3 *P*=9.8 ×10^−4^; C4 *P*=0.0068; T1 *P*=2.2×10^−5^).

### TDP-43 HISTOPATHOLOGY IN PRIMARY MOTOR CORTEX

Staining for unphosphorylated TDP-43 in M1 of a control monkey (monkey W) revealed widespread nuclear staining as expected, but also a band of light-staining in both the nucleus and cytoplasm of the large Betz cells of layer Vb (Fig. 6A). This suggests that these specialized cells may have a high level of functional TDP-43 expression even in healthy animals. In the transfected animals, TDP-43 staining in Betz cells was more dense (Fig. 6B, C).

**Figure 6.**
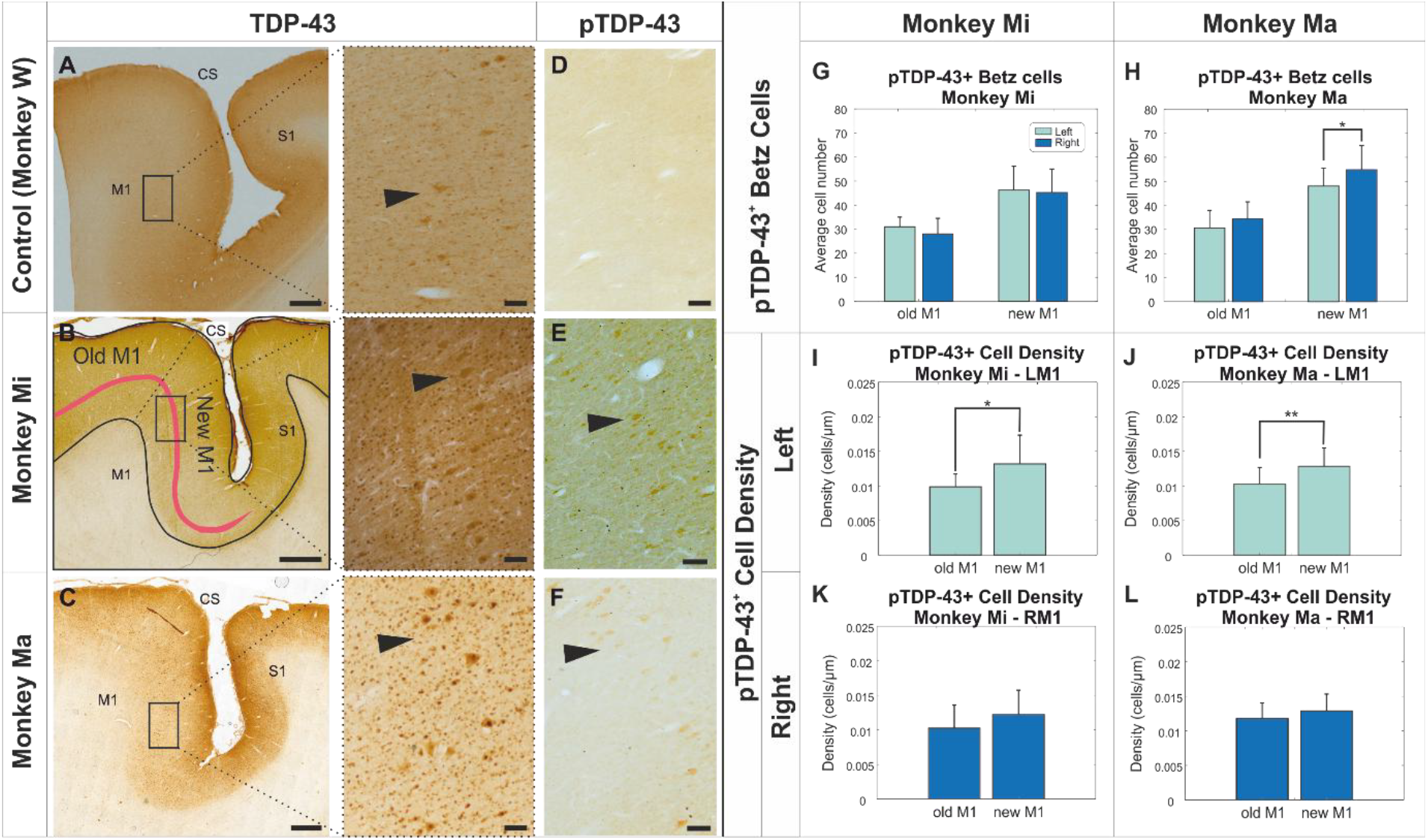
TDP-43 and pTDP-43 in the sensorimotor cortex. (A-F) Images show parasagittal sections of the cortex in the region surrounding the central sulcus (CS). Pre-central (primary motor cortex, M1) and post-central (primary somatosensory cortex, S1) cortices are labelled. Sections have been stained to label TDP-43 or pTDP-43 in brown, using DAB. (**A**) TDP-43 was found in the cytoplasm of the large Betz cells of layer V (indicated by arrows) in M1 of a healthy control macaque (Monkey W), without virus transfection (see enlarged view). (**B, C**) In the transfected animals, Monkey Mi and Monkey Ma, staining was more dense, with clear cytoplasmic staining of Betz cell cytoplasm and nuclei in the enlarged images. (**D**) No pTDP-43 expression was detected in M1 of the healthy control macaque (Monkey W). **(E, F)** Staining for pTDP-43 in the transfected animals identified dense staining in the cytoplasm of layer Vb cells (indicated by arrows). Nuclear staining was present but less prominent. **(G, H)** Counts of pTDP-43+ Betz cells in two subdivisions of M1, in two transfected animals. Examples of the division of New M1 and Old M1 are shown in (B). (**L-O**) the density of pTDP-43+ cell was calculated as cells per unit length of layer Vb. Plots show results for left and right M1, for each monkey separately. Scale bars: low power images: 1000µm; high power images: 100µm. Bars in (G-L) are the average over 15-22 sections sampled from the mediolateral extent of M1; error bars indicate standard error of the means. Asterisks indicate significant differences: *, P<0.05; **, P<0.005.

No staining for the pathological pTDP-43 was seen in the control monkey (Fig. 6D), whereas it was clearly seen in the cytoplasm, but not the nucleus of the Betz cells in the transfected animals (Fig. 6E, F). The expression appeared uniform throughout the cytoplasm and did not show any signs of aggregation or clumping. This is consistent with the report that TDP-43 proteinopathy in Betz cells differs from that in spinal and bulbar motor neurons[41].

In primates, M1 can be subdivided into New M1, which lies within the bank of the central sulcus and has many fast direct cortico-motoneuronal connections, and Old M1, which is situated on the cortical surface and makes indirect, as well as slower direct connections to motor neurons[35, 36]. We quantified the distribution of pTDP-43+ Betz cells within these two divisions as indicated on Fig. 6B. Previously described anatomical markers[35] were used to define the boundary between Old and New M1; the outer borders were defined by the lack of giant Betz cells. A higher number of pTDP-43+ Betz cells were seen in New M1 than Old M1 in both transfected animals (Fig. 6G, H). Significantly more pTDP-43+ Betz cells were seen in right compared to left New M1 in Monkey Ma (*P*=0.0188, Fig. 6H), but no side-to-side differences were seen in Monkey Mi (Fig. 6G).

To account for differences in size of New and Old M1, we calculated the density of pTDP-43+ cells by dividing the raw counts by the linear extent of layer Vb in each area. Labelled cell density was significantly higher in New M1 than in Old M1 for the left hemisphere in both animals (Monkey Mi, P=0.0359; Monkey Ma, P=0.0037). No such differences between New and Old M1 were seen in the right hemisphere (Fig. 6I-L).

## DISCUSSION

### FOCAL OVEREXPRESSION OF TDP-43 PRODUCES WIDESPREAD NEUROPATHOLOGY

A critical question in motor neuron disease research is the role of TDP-43 pathology. While mislocalized TDP-43 aggregates are a consistent histopathological hallmark observed in the tissue from almost all patients, it remains unclear whether this plays a causal role in the initiation and spreading of pathology. The alternative, that TDP-43 abnormalities are a (maladaptive or adaptive) response to disease progression remains a viable possibility. Here we addressed this issue experimentally by producing highly focal overexpression of TDP-43 in a single motor neuron pool using an intersectional genetics approach. Whatever the role of TDP-43 in human disease, we might expect that powerful overexpression of a single protein would be toxic to healthy cell function. It is unsurprising that such transfected cells should eventually die, as appears to have happened in our model (Fig. 4E). However, if TDP-43 is a purely downstream effect of degeneration, pathological changes should be confined to the initially targeted cell population. Contrary to this, we observed widespread TDP-43 pathology extending throughout the spinal cord and reaching the motor cortex. These findings support the hypothesis that TDP-43 dysfunction itself is involved in the dissemination of neuropathology beyond the site of initial insult, implicating it as an active contributor to disease progression.

The extent of TDP-43 pathology observed here was remarkable. High levels of pTDP-43 were seen in motor neurons and in smaller cells outside Rexed’s lamina IX (presumed interneurons) on both sides of the spinal cord, and over a wide range of spinal segments (Fig. 5I, J). In M1, pTDP-43 was present in both Old M1 and New M1 bilaterally and over a wide medio-lateral extent (Fig. 6G-L). Spread of pTDP-43 was not therefore confined to cells directly connected to the transfected motor neurons innervating the brachioradialis muscle. This is likely to reflect multiple waves of disease spread, in which successively more indirectly connected neurons were recruited to the pathological process. By the time the animals were killed and tissue examined, this process seems to have become well advanced, and little signature of the initial overexpression site remained. However, it was notable that in the cortex of both monkeys, a higher density of pTDP-43+ Betz cells were seen in left New M1 than Old M1 (Fig. 6I, J); left New M1 has the most direct connections to the motor neurons of the right cord where initial overexpression occurred. Due to the advanced stage of neuropathological spread, we can provide limited information on the route of that spread. For example, spread to the ipsilateral intermediate zone and to the contralateral spinal cord could equally have occurred via intra-spinal connections, or passed via the cortex. However, since in our model the initial overexpression was confined to lower motor neurons, the presence of pTDP-43 in the cortex implies that retrograde spread is possible, consistent with the ‘dying backward’ hypothesis. Our results do not exclude the possibility of anterograde spread, from cortex to spinal cord, as posited by the ‘dying forward’ model.

As previously reported, we found that TDP-43 was restricted to the nucleus in healthy spinal tissue. Surprisingly, in healthy macaque M1 this was not the case: the cytoplasm of large layer Vb Betz cells stained lightly for TDP-43. This raises the possibility that these unusual corticospinal cells, with their long, fast conducting axons, extensive terminal arborization and membrane conductance specialized for fast discharge rates, may be uniquely vulnerable to pathological phosphorylation of TDP-43 since they have higher levels of functional protein available.

Research in non-human primates necessarily places ethical and financial limits on the number of animals which can be used. In this study, this meant that we were only able to examine tissue at a single time point 32 weeks after virus injection. At this time, although both animals showed extensive pTDP-43 neuropathology, there was no evidence of widespread motor neuron degeneration. The animals showed no sign of clinical weakness, and there were no further regions of hyperintensity on the MRI scan after the initial transient signals starting 9-15 weeks after viral injection. It is possible that the experiment was ended before the accumulation and cytoplasmic mislocalization of pathological TDP-43 outside the brachioradialis motor neuron pool became sufficiently toxic to produce cell death. Alternatively, further degeneration may have occurred, but at a steady low rate which was subthreshold for detection on MRI. In humans, motor unit number estimation techniques identify deficits prior to clinical presentation of disease[42, 43], and it is estimated that around 20% of motor neurons are lost before patients first complain of weakness[44]; we would presumably also be unlikely to detect overt weakness in monkeys before many motor neurons have been lost. This model allows us the unique opportunity to study the early phase of disease, and so better understand the initial pathological propagation of TDP-43.

It is important to consider possible confounding factors which may have led to the widespread TDP-43 pathology. With our intersectional genetics approach, viral transfection with AAV9 was intentionally widespread throughout the cord. Viral transfection of rAAV2 relied upon retrograde transport and thus was confined to the motor neurons supplying the brachioradialis muscle. In this way, viral transfection and thus, transduction and overexpression of TDP-43 would be restricted to only the motor pool of the right brachioradialis muscle, providing a focal site of onset. Several lines of evidence support that this was indeed the case. Firstly, an extensive literature has reported focal retrograde transport and expression following intramuscular injections of viral vectors in primates[45, 46], which differs from the extensive CNS transfection reported after intravenous administration in primates[47-49] and after intramuscular administration in neonatal mice[50]. Secondly, some 8-14 weeks after the initial viral injection, focal changes became apparent in the right brachioradialis muscle on MRI scans. There were no more diffuse changes at this time, as would be expected if the virus had spread more widely. Finally, any TDP-43 produced from overexpression following viral transfection should be accompanied by the fluorescent tag tdTomato, which was incorporated into the genetic payload in the rAAV2 virus. In prior viral delivery experiments, we reliably detected tdTomato labelled cells, confirming successful transduction and transgene expression (Fig. 4A). However, in the two animals examined in this study, no tdTomato labelled cells were identified at the time of *post-mortem* analysis. Given our previous success using the same construct and detection methods, the absence of tdTomato signal in these animals may indicate that all such transfected cells had died by the time the tissue was examined *post-mortem*. This interpretation is consistent with the reduction in ChAT positive cells in the right side of cervical segments which innervate the injected muscle, as well as the bands of signal intensity detected in the right brachioradialis on MRI. The absence of tdTomato labelled cells in other regions of the spinal cord or motor cortex further supports the conclusion that the rAAV2 vector did not spread systemically via the bloodstream, consistent with previous reports of focal transfection following intramuscular delivery in adult primates[45, 46].

### A PRIMATE MODEL OF MOTOR NEURON DISEASE

An important aspect of this study was the use of macaque monkeys, an Old World primate with very similar motor system to humans. Macaques have key differences from the rodent species more commonly used as a model of motor neuron disease[51]. The primate corticospinal tract runs primarily in the dorsolateral funiculus, compared to the dorsal column in rats and mice. Some corticospinal neurons in primates make monosynaptic connections to lower motor neurons, a connection which is important for the performance of independent finger movements[37]. These connections originate mainly in the subdivision of M1 referred to as New M1 by Strick and co-workers[35, 52] (although not exclusively[36]), which is unique to primates. In rodents, the corticospinal tract connects primarily to dorsal horn interneurons, involved in sensory processing; in primates most connections are by contrast to the intermediate zone, providing indirect control of motor neuron firing[37]. These differences may be important in how motor neuron disease spreads trans-synaptically, especially between the cortex and the spinal cord. In our data, the density of pTDP-43+ cells was higher in New M1 than Old M1 in both monkeys on the side contralateral to the initially transfected brachioradialis motor neuron pool. Since New M1 is the main location of fast cortico-motoneuronal cells, this suggests that these uniquely primate connections may be of importance for disease spread.

In both rodents and primates, the majority of corticospinal axons are slowly conducting; however, primates also have a small number of cells with fast conduction velocities >50 m/s[53]. These cells express the fast potassium channel Kv3.1b, leading to narrow action potentials[54]. In rodents, Kv3.1b is seen only in fast-spiking interneurons not in pyramidal cells. This is likely to be just one of the specialist adaptations of primate Betz cells to permit high discharge rates and fast axonal conduction velocities. Interestingly, in control tissue from a healthy macaque we observed light cytoplasmic staining for TDP-43 in the Betz cells of the motor cortex (Fig. 6A), which is not seen in healthy rodents[55]. It is possible that this finding is another reflection of the functional specialization and high energy turnover of these cells. This may make these upper motor neurons especially vulnerable to pathological protein aggregation, since they have a ready store of cytoplasmic functional TDP-43 which can be converted to pathological pTDP-43.

The function and conformation of TDP-43 is highly conserved between humans, rodents and eukaryotes[56, 57]; however, recent reports have identified key differences in its role in mRNA regulation between species. Recent advances have identified an important role for TDP-43 in the repression of non-conserved cryptic exons, with loss of nuclear TDP-43 associated with mRNA splicing defects and anomalous translation [58]. Stathmin-2 (STMN2) mRNA, which encodes a protein essential for axonal growth and repair in neurons, is one of many mRNAs found to be regulated by TDP-43 [59-61]. The loss or altered functioning of TDP-43 in human motor neurons has been shown to induce aberrant splicing and polyadenylation of stathmin-2 mRNA, causing a significant reduction in levels of functional mRNA [62, 63]. Aberrant alterations of stathmin-2 mRNA in motor neurons of ALS patients, as well as the ability of the functional protein to rescue the disease phenotype have implicated the TDP-43-stathmin-2 interaction in motor neuron degeneration. Stathmin-2 shows sequence homology between humans, Old World and some New World primates, but not rodents[63]. A model such as ours would enable the study of these conserved downstream molecular components at various stages of disease presentation.

### MUSCLE MRI FOR DETECTION OF MOTOR NEURON DEGENERATION

Structural MRI scans have long been used to assess cortical thinning and, more recently, structural differences in the motor cortices have been used to distinguish ALS patients from healthy controls[64, 65]. In addition to these cortical assessments, scans of muscle could reveal on-going denervation in ALS patients and have potential as a disease biomarker[66]. T2-weighted RARE scans detect fluctuations in the proportion of water within a muscle. Following denervation, muscle cytoarchitecture becomes destabilized, leading to hyperintense signal[26]. In this study we detected bands of high signal in the transfected brachioradialis muscle, eight and fourteen weeks after the virus injection in the two respective monkeys, which increased and spread through the muscle before recovering. This increase occurred too late to be associated merely with inflammation caused by the intramuscular virus injections, and is most likely explained by denervation caused by TDP-43 overexpression in transfected motor neurons. The recovery of this signal after around eight weeks is presumably explained by compensatory reinnervation and motor unit remodeling.

In monkey Mi, hyperintense signal from the brachioradialis muscle on MRI began eight weeks after injection, compared with fourteen weeks after injection for monkey Ma. This suggests that overexpression of TDP-43 may have reached toxic levels earlier in monkey Mi than Ma. At *post-mortem*, monkey Ma showed a characteristic peak in motor neuron number around the cervical enlargement (C6-C8), similar to that seen in the control animal (Fig. 4D, E). By contrast, in monkey Mi this peak was abolished, which may reflect the onset of secondary degeneration of motor neurons close to, but not part of, the transfected brachioradialis motor neuron pool following the earlier disease spread in this animal.

While the time of onset of the hyperintense MRI signal differed between the two animals, the duration and spatial progression throughout the muscle was conserved. No further indication of denervation was recorded in either the virus-targeted muscle or any of the other forearm muscles. Whilst the initial denervation event provoked widespread cytoarchitectonic destabilization throughout the right brachioradialis, later more gradual reorganization caused by progressive spread may not have generated sufficiently extensive or coordinated changes that could be resolved with this technique. Alternatively, denervation in other forearm muscles may not have been detected on MRI as the experiment may have been terminated before spread of pTDP-43 reached levels sufficiently toxic to induce motor neuron cell death in these motor pools.

### OUTLOOK

In this report we have described the development of a non-human primate model of amyotrophic lateral sclerosis, which replicates the progressive spread of TDP-43 pathology seen in humans. Importantly, non-invasive MRI scans allowed us to monitor the onset of the initial transfection-induced degeneration event. This occurred at variable times after virus injection in the two monkeys; pinpointing degeneration onset in this way could allow subsequent experimental disease-modifying therapies to be initiated with accurate timing. This model has great potential for pre-clinical testing; the use of a primate species with close similarities to humans increases the likelihood of promising results translating to clinical benefit. In addition, the availability of such a model could allow the testing and validation of early biomarkers[67], which is not possible in patients who are typically diagnosed quite late in the disease time course.

## RESOURCE AVAILABILITY

### Lead contact

Further information and requests for resources and reagents should be directed to and will be fulfilled by the lead contact, Prof Stuart Baker.

### Data and code availability

Raw MRI data, and spreadsheets containing cell counts, will be made publicly available on the server https://data.ncl.ac.uk after acceptance of this paper.

## ACKNOWLEDGMENTS

The authors would like to thank Terri Jackson and Andrew Atkinson for assistance with behavioral training, Kathy Murphy and Rocio Fernandez-Palacios O’Connor for veterinary and anesthesia support, Ashley Waddle and his team for animal care, Norman Charlton and Nathan Cassie for mechanical engineering, and Michelle Waddle for theatre assistance. We also wish to thank Richard Leech and the late Roy Leech, whose enthusiasm for this research led to the project’s inception.

This work was funded by The William Leech Charity.

## AUTHOR CONTRIBUTIONS

Conceptualization, M.R.B and S.N.B.; methodology, R.H.A.J, M.R.B. and S.N.B.; Investigation, R.H.A.J., L.S., F.B., I.S. and C.M.; writing—original draft, R.H.A.J, M.R.B. and S.N.B.; writing—review & editing, R.H.A.J, M.R.B. and S.N.B.; funding acquisition, M.R.B. and S.N.B.; resources, M.R.B. and S.N.B.; supervision, M.R.B. and S.N.B.

## REFERENCES

1. Ragagnin, A.M.G., et al., Motor Neuron Susceptibility in ALS/FTD. Front Neurosci, 2019. 13: p. 532.

2. Marin, B., et al., Variation in worldwide incidence of amyotrophic lateral sclerosis: a meta-analysis. Int J Epidemiol, 2017. 46(1): p. 57–74.

3. Gao, J., et al., Pathomechanisms of TDP-43 in neurodegeneration. J Neurochem, 2018.

4. Buratti, E. and F.E. Baralle, Multiple roles of TDP-43 in gene expression, splicing regulation, and human disease. Frontiers in Bioscience-Landmark, 2008. 13: p. 867–878.

5. Arai, T., et al., TDP-43 is a component of ubiquitin-positive tau-negative inclusions in frontotemporal lobar degeneration and amyotrophic lateral sclerosis. Biochem Biophys Res Commun, 2006. 351(3): p. 602–11.

6. Neumann, M., et al., Ubiquitinated TDP-43 in frontotemporal lobar degeneration and amyotrophic lateral sclerosis. Science, 2006. 314(5796): p. 130–3.

7. Brettschneider, J., et al., Stages of pTDP-43 Pathology in Amyotrophic Lateral Sclerosis. Annals of Neurology, 2013. 74(1): p. 20–38.

8. Ravits, J.M. and A.R. La Spada, ALS motor phenotype heterogeneity, focality, and spread: deconstructing motor neuron degeneration. Neurology, 2009. 73(10): p. 805–11.

9. Weber, M., et al., The split hand in ALS has a cortical basis. Journal of the Neurological Sciences, 2000. 180(1-2): p. 66–70.

10. Fallini, C., G.J. Bassell, and W. Rossoll, The ALS disease protein TDP-43 is actively transported in motor neuron axons and regulates axon outgrowth. Hum Mol Genet, 2012. 21(16): p. 3703–18.

11. Braak, H., et al., Amyotrophic lateral sclerosis-a model of corticofugal axonal spread. Nature Reviews Neurology, 2013. 9(12): p. 708–714.

12. Eisen, A., et al., Cortical influences drive amyotrophic lateral sclerosis. Journal of Neurology Neurosurgery and Psychiatry, 2017. 88(11): p. 917–924.

13. Iwatsubo, T., et al., Corticofugal projections to the motor nuclei of the brainstem and spinal cord in humans. Neurology, 1990. 40(2): p. 309–12.

14. Horn, A.K. and R.J. Leigh, The anatomy and physiology of the ocular motor system. Handb Clin Neurol, 2011. 102: p. 21–69.

15. Kiernan, J.A. and A.J. Hudson, Changes in Sizes of Cortical and Lower Motor Neurons in Amyotrophic-Lateral-Sclerosis. Brain, 1991. 114: p. 843–853.

16. Dadon-Nachum, M., E. Melamed, and D. Offen, The “Dying-Back” Phenomenon of Motor Neurons in ALS. Journal of Molecular Neuroscience, 2011. 43(3): p. 470–477.

17. Martineau, E., et al., Dynamic neuromuscular remodeling precedes motor-unit loss in a mouse model of ALS. Elife, 2018. 7.

18. Baker, M.R., ALS-dying forward, backward or outward? Nature Reviews Neurology, 2014. 10(11).

19. Rosen, D.R., Mutations in Cu/Zn Superoxide-Dismutase Gene Are Associated with Familial Amyotrophic-Lateral-Sclerosis (Vol 362, Pg 59, 1993). Nature, 1993. 364(6435): p. 362–362.

20. Deng, H.X., et al., Conversion to the amyotrophic lateral sclerosis phenotype is associated with intermolecular linked insoluble aggregates of SOD1 in mitochondria. Proceedings of the National Academy of Sciences of the United States of America, 2006. 103(18): p. 7142–7147.

21. Ozdinler, P.H., et al., Corticospinal motor neurons and related subcerebral projection neurons undergo early and specific neurodegeneration in hSOD1G(9)(3)A transgenic ALS mice. J Neurosci, 2011. 31(11): p. 4166–77.

22. Heffner, R. and B. Masterton, Variation in form of the pyramidal tract and its relationship to digital dexterity. Brain Behav Evol, 1975. 12(3): p. 161–200.

23. Uchida, A., et al., Non-human primate model of amyotrophic lateral sclerosis with cytoplasmic mislocalization of TDP-43. Brain, 2012. 135(Pt 3): p. 833–46.

24. Jones, R.H.A., A macaque model of Motor Neurone Disease. Newcastle University, 2022. Ph. D Thesis. Available at: http://theses.ncl.ac.uk/jspui/handle/10443/5749.

25. Kullmer, K., et al., Changes of sonographic, magnetic resonance tomographic, electromyographic, and histopathologic findings within a 2-month period of examinations after experimental muscle denervation. Archives of Orthopaedic and Trauma Surgery, 1998. 117(4-5): p. 228–234.

26. Kamath, S., et al., MRI appearance of muscle denervation. Skeletal Radiol, 2008. 37(5): p. 397–404.

27. Viddeleer, A.R., et al., Quantitative STIR of muscle for monitoring nerve regeneration. J Magn Reson Imaging, 2016. 44(2): p. 401–10.

28. Schneider, C.A., W.S. Rasband, and K.W. Eliceiri, NIH Image to ImageJ: 25 years of image analysis. Nat Methods, 2012. 9(7): p. 671–5.

29. Tournier, J.D., et al., MRtrix3: A fast, flexible and open software framework for medical image processing and visualisation. Neuroimage, 2019. 202: p. 116137.

30. Benjamini, Y. and Y. Hochberg, Controlling the False Discovery Rate - a Practical and Powerful Approach to Multiple Testing. Journal of the Royal Statistical Society Series B-Statistical Methodology, 1995. 57(1): p. 289–300.

31. Meyer, K., et al., Improving Single Injection CSF Delivery of AAV9-mediated Gene Therapy for SMA: A Dose-response Study in Mice and Nonhuman Primates. Molecular Therapy, 2015. 23(3): p. 477–487.

32. Schnell, S.A., W.A. Staines, and M.W. Wessendorf, Reduction of lipofuscin-like autofluorescence in fluorescently labeled tissue. Journal of Histochemistry & Cytochemistry, 1999. 47(6): p. 719–730.

33. Inukai, Y., et al., Abnormal phosphorylation of Ser409/410 of TDP-43 in FTLD-U and ALS. FEBS Lett, 2008. 582(19): p. 2899–904.

34. O’Brien, J., H. Hayder, and C. Peng, Automated Quantification and Analysis of Cell Counting Procedures Using ImageJ Plugins. Jove-Journal of Visualized Experiments, 2016(117).

35. Rathelot, J.A. and P.L. Strick, Subdivisions of primary motor cortex based on cortico-motoneuronal cells. Proceedings of the National Academy of Sciences of the United States of America, 2009. 106(3): p. 918–923.

36. Witham, C.L., et al., Corticospinal Inputs to Primate Motoneurons Innervating the Forelimb from Two Divisions of Primary Motor Cortex and Area 3a. Journal of Neuroscience, 2016. 36(9): p. 2605–2616.

37. Kuypers, H.G.J.M. 1981, ‘Anatomy of the descending pathways.’ In: Brooks, V., Ed., The Nervous System, Handbook of Physiology, Vol 2, Williams and Wilkins, Baltimore, 597–666.

38. Dum, R.P. and P.L. Strick, Spinal cord terminations of the medial wall motor areas in macaque monkeys. Journal of Neuroscience, 1996. 16(20): p. 6513–6525.

39. Jenny, A.B. and J. Inukai, Principles of motor organization of the monkey cervical spinal cord. J Neurosci, 1983. 3(3): p. 567–75.

40. Moreno-Garcia, A., et al., An Overview of the Role of Lipofuscin in Age-Related Neurodegeneration. Frontiers in Neuroscience, 2018. 12.

41. Braak, H., et al., Pathological TDP-43 changes in Betz cells differ from those in bulbar and spinal alpha-motoneurons in sporadic amyotrophic lateral sclerosis. Acta Neuropathol, 2017. 133(1): p. 79–90.

42. Escorcio-Bezerra, M.L., et al., Motor Unit Number Index and Neurophysiological Index as Candidate Biomarkers of Presymptomatic Motor Neuron Loss in Amyotrophic Lateral Sclerosis. Muscle & Nerve, 2018. 58(2): p. 204–212.

43. Neuwirth, C., et al., Motor Unit Number Index (MUNIX) detects motor neuron loss in pre-symptomatic muscles in Amyotrophic Lateral Sclerosis. Clinical Neurophysiology, 2017. 128(3): p. 495–500.

44. Aggarwal, A. and G. Nicholson, Detection of preclinical motor neurone loss in SOD1 mutation carriers using motor unit number estimation. Journal of Neurology Neurosurgery and Psychiatry, 2002. 73(2): p. 199–201.

45. Towne, C., et al., Efficient transduction of non-human primate motor neurons after intramuscular delivery of recombinant AAV serotype 6. Gene Ther, 2010. 17(1): p. 141–6.

46. Bohlen, M.O., H.G. El-Nahal, and M.A. Sommer, Transduction of Craniofacial Motoneurons Following Intramuscular Injections of Canine Adenovirus Type-2 (CAV-2) in Rhesus Macaques. Frontiers in Neuroanatomy, 2019. 13.

47. Samaranch, L., et al., Adeno-associated virus serotype 9 transduction in the central nervous system of nonhuman primates. Hum Gene Ther, 2012. 23(4): p. 382–9.

48. Jackson, K.L., et al., Initial gene vector dosing for studying symptomatology of amyotrophic lateral sclerosis in non-human primates. J Med Primatol, 2015. 44(2): p. 66–75.

49. Bevan, A.K., et al., Systemic gene delivery in large species for targeting spinal cord, brain, and peripheral tissues for pediatric disorders. Mol Ther, 2011. 19(11): p. 1971–80.

50. Chen, Z., et al., rAAV2-Retro Enables Extensive and High-Efficient Transduction of Lower Motor Neurons following Intramuscular Injection. Mol Ther Methods Clin Dev, 2020. 17: p. 21–33.

51. Lemon, R.N. and J. Griffiths, Comparing the function of the corticospinal system in different species: organizational differences for motor specialization? Muscle Nerve, 2005. 32(3): p. 261–79.

52. Rathelot, J.A. and P.L. Strick, Muscle representation in the macaque motor cortex: An anatomical perspective. Proceedings of the National Academy of Sciences of the United States of America, 2006. 103(21): p. 8257–8262.

53. Firmin, L., et al., Axon diameters and conduction velocities in the macaque pyramidal tract. J Neurophysiol, 2014. 112(6): p. 1229–40.

54. Soares, D., et al., Expression of Kv3.1b potassium channel is widespread in macaque motor cortex pyramidal cells: A histological comparison between rat and macaque. Journal of Comparative Neurology, 2017. 525(9): p. 2164–2174.

55. Ding, X.B., et al., Spreading of TDP-43 pathology via pyramidal tract induces ALS-like phenotypes in TDP-43 transgenic mice. Acta Neuropathologica Communications, 2021. 9(1).

56. Wang, H.Y., et al., Structural diversity and functional implications of the eukaryotic TDP gene family. Genomics, 2004. 83(1): p. 130–139.

57. Ayala, Y.M., et al., Human, Drosophila, and C-elegans TDP43: Nucleic acid binding properties and splicing regulatory function. Journal of Molecular Biology, 2005. 348(3): p. 575–588.

58. Ling, J.P., et al., TDP-43 repression of nonconserved cryptic exons is compromised in ALS-FTD. Science, 2015. 349(6248): p. 650–5.

59. Chauvin, S. and A. Sobel, Neuronal stathmins: a family of phosphoproteins cooperating for neuronal development, plasticity and regeneration. Prog Neurobiol, 2015. 126: p. 1–18.

60. Shin, J.E., S. Geisler, and A. DiAntonio, Dynamic regulation of SCG10 in regenerating axons after injury. Exp Neurol, 2014. 252: p. 1–11.

61. Shin, J.E., et al., SCG10 is a JNK target in the axonal degeneration pathway. Proc Natl Acad Sci U S A, 2012. 109(52): p. E3696–705.

62. Klim, J.R., et al., ALS-implicated protein TDP-43 sustains levels of STMN2, a mediator of motor neuron growth and repair. Nat Neurosci, 2019. 22(2): p. 167–179.

63. Melamed, Z., et al., Premature polyadenylation-mediated loss of stathmin-2 is a hallmark of TDP-43-dependent neurodegeneration. Nat Neurosci, 2019. 22(2): p. 180–190.

64. Roccatagliata, L., et al., Detection of motor cortex thinning and corticospinal tract involvement by quantitative MRI in amyotrophic lateral sclerosis. Amyotrophic Lateral Sclerosis, 2009. 10(1): p. 47–52.

65. Ferraro, P.M., et al., Multimodal structural MRI in the diagnosis of motor neuron diseases. Neuroimage Clin, 2017. 16: p. 240–247.

66. Jenkins, T.M., et al., Longitudinal multi-modal muscle-based biomarker assessment in motor neuron disease. Journal of Neurology, 2020. 267(1): p. 244–256.

67. Poesen, K. and P. Van Damme, Diagnostic and Prognostic Performance of Neurofilaments in ALS. Frontiers in Neurology, 2019. 9.

